# Investigating the growth of an engineered strain of Cyanobacteria with an Agent-Based Model and a Recurrent Neural Network

**DOI:** 10.1101/2021.10.11.463942

**Authors:** Jonathan Sakkos, Joe Weaver, Connor Robertson, Bowen Li, Denis Taniguchi, Ketan Maheshwari, Daniel Ducat, Paolo Zuliani, Andrew Stephen McGough, Tom Curtis, Miguel Fuentes-Cabrera

## Abstract

A computational framework combining Agent-Based Models (ABMs) and Deep Learning techniques was developed to help design microbial communities that convert light and CO_2_ into useful bioproducts. An ABM that accounts for CO_2_, light, sucrose export rate and cell-to-cell mechanical interactions was used to investigate the growth of an engineered sucrose-exporting strain of *Synechococcus elongatus* PCC 7942. The ABM simulations produced population curves and synthetic images of colony growth. The curves and the images were analyzed, and growth was correlated to nutrients availability and colonies’ initial spatial distribution. To speed up the ABM simulations, a metamodel based on a Recurrent Neural Network, RNN, was trained on the synthetic images of growth. This metamodel successfully reproduced the population curves and the images of growth at a lower computational cost. The computational framework presented here paves the road towards designing microbial communities containing sucrose-exporting *Synechococcus elongatus* PCC 7942 by exploring the solution space *in silico* first.

## 1 Introduction

Microbial communities contribute important functions towards human health and biotechnology, including the many microbes found associated with the human body (*e.g.*, gut microbiome) and species that contribute to soil fertility and plant health. Many important features of microbial communities (*e.g.*, robustness, production of target compounds, efficient utilization of resources) arise as emergent properties from interactions between individual species. The increasing recognition of the ubiquity and importance of natural communities has spurred interest in designer microbial consortia which might be tailored for a range of useful outputs, while retaining important emergent features of their natural counterparts [1]. Yet, the increased complexity of mixed species consortia creates additional difficulties in engineering optimal configurations, both in the technical capabilities of modifying more than one strain as well as in accurate prediction of the impact of the changes on communal output. The increased complexity of mixed-species communities and lack of systems-level predictive tools are currently substantial barriers to their adoption for bioindustrial applications. Development of versatile platforms that can be used to construct and evaluate a range of simple mixed microbial communities could enable bottom up strategies to better understand and design complex microbial consortia [2].

One previously-developed biological platform for flexibly constructing microbial co-cultures involves an engineered strain of the model cyanobacterium, *Synechococcus elongatus* PCC 7942, that exports large quantities of sucrose. The secreted cyanobacterial sucrose can directly support co-cultivated heterotrophic microbes and this system has been used to build a number of co-cultures that are stable over long time periods utilizing different heterotrophic species [3, 4, 5, 6, 7, 8, 9]. Such communities have been programmed to convert light and CO_2_ into a variety of useful bioproducts (*e.g.*, fuels or bioplastics) by utilizing the heterotrophic species to convert cyanobacterial sucrose into other metabolites. Nonetheless, designing and optimizing this type of communities remains a challenge, largely due to the number of parameters to vary, the cost and time to perform pure wet-lab experiments exploring the solution space, and the lack of predictive models used to guide them.

Agent-Based Models, ABMs, of microbial growth can be a useful tool for understanding such communities. These models allows exploration of the solution space *in silico*, thus potentially save considerable experimental time and expense. In recent years there has been many, and significant, successes in the applications of ABM techniques to microbiology, see [10, 11, 12] and references therein. Unfortunately, two major roadblocks still limit the applicability of ABM in this field: i) simulating communities which exhibit spatial and temporal dimensions that resemble those of real experiments; ii) gauging the accuracy of the ABM. Neither of these roadblocks are easy to overcome. The computational cost of running simulations in microbiology at the scale required to approach those of real experiments can be astronomical, especially as the execution time can grow exponentially with the number of agents and their interactions. The computational cost is compounded by the fact that for gauging the accuracy of ABMs, it’s recommend to use an approach known as *Pattern Oriented Modeling*, POM, in which the patterns produced by the simulation and the experiment at many different temporal and spatial scales are compared[13].

We hypothesize that an approach that combines ABMs that run in High-Performance Computers, HPCs, with metamodels built with Deep Learning techniques, DL, could overcome the aforementioned roadblocks. Here, we took the first steps to probe this hypothesis. For this, we investigated the growth of a microbial community made from the sucrose exporting cyanobacteria [3] using NUFEB [14], and a metamodel thereof based on a Recurrent Neural Network, RNN. NUFEB is a massively parallel ABM simulator capable of investigating microbial communities with up to 10^7^ microbes. Here, however, we used NUFEB to investigate a community with ca up to 10^5^ microbes. The data produced by the simulations included the positions of individual cells and colonies during growth, and the population curves of the whole microbial community. This data was subsequently analyzed and metrics that relate growth to the spatial patterns of individual colonies and nutrients availability were extracted. An RNN is an artificial neural network that is designed to treat data represented by a time-series. The RNN learns the spatial-temporal patterns in the series and when trained appropriately can make accurate predictions. Here we investigated whether a metamodel based on an RNN could be used to accurately predict the growth of the simulated microbial community, reducing the computational cost associated with running our ABM. Specifically, we used the synthetic images of colony growth generated by the ABM to train predRNN [15], which was subsequently used to make predictions of microbial growth.

## 2 Method

### 2.1 Agent Based Model

An agent based representation of the sucrose export cyanobacteria [3] (for simplicity, from now on we will refer to this bacteria as cyanobacteria) was developed and integrated into NUFEB [14] for studying its spatial and temporal dynamics. In the ABM, the computational domain is defined as the area where cyanobacteria reside and where the biological, physical and chemical processes take place. This area is a 3D cuboid with dimensions 100*μm ×* 100*μm ×* 10*μm*. Within this area, nutrients are represented as continuous field where their dynamics over time and space is updated at each discrete Cartesian grid. Cyanobacteria cells are modelled as spherical particles with each having a set of state variables to describe its physical and biological attributes (position, size, growth rate, etc.) These attributes vary between cells and can change over time as a result of external or internal processes. The processes that influence cyanobacteria activities are classified into three sub-models: biological, physical, and chemical.

The *biological sub-model* handles metabolism, growth, and reproduction of cyanobacteria. An individual cell grows and its mass increase by consuming nutrients in the grid where it is located. The equation governing the mass *m*_*i*_ of cyanobacteria *i* is given by

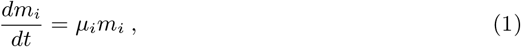

where a Monod-based growth model is implemented to determine the specified growth rate *μ*, which depends on CO_2_ and light

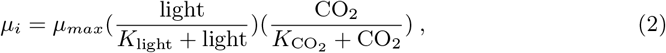

where light and CO_2_ are the amounts of light and CO_2_ present around the cell. *μ*_*max*_ is the maximum growth rate, *K*_light_ and 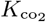 are affinity constants for light and CO_2_, respectively. The values of *μ*_*max*_, *K*_light_ and 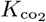 are obtained from experimental data[16, 17].

Bacteria division is the result of cell growth, which is considered as an instantaneous process in the model. Cells are modelled as spheres within NUFEB [14] and since cyanobacterium is rod shaped, the volume-equivalent diameter was used to more accurately characterize the cell size and biomass. It is assumed that the division occurs when the diameter of a cyanobacterium reaches 1.96*μm*. The cell then divides into two daughter cells, one of which takes the location of the parent cell while the centre of the other cell is randomly chosen at a distance corresponding to the sum of the diameter of the daughters. The total mass of the two daughter cells is always conserved from the parent cell. During the division, one daughter has a mass (uniformly) randomly chosen between 40% − 60% of the parent cell’s mass, with the other cell inheriting the remainder.

Mechanical interactions between cyanobacteria are handled by the *physical sub-model*, which is built upon our previous mechanical ABM [18]. When cells grow and divide, the system can deviate from mechanical equilibrium (*i.e.*, non-zero net force on cells) due to cell overlap and collision. Mechanical relaxation is therefore required to update the positions of the cells and minimize the stored mechanical energy of the system. More specifically, the net force acting on an agent is calculated as the sum of frictional and viscous damping forces. The former is imposed to push two agents apart when the distance between them is less than their contact distance (*i.e.*, the sum of their diameters) based on Hooke’s law, as described in [19]. The latter can be thought of as a resistance drag force caused by the motion of spherical particles in a fluidic environment. The relaxation is carried out using the discrete element method, and the Newtonian equations of motion are solved by using Verlet algorithm provided by the software LAMMPS [20].

The processes included in the *chemical sub-model* aim to update nutrient consumption and their mass balance within the grid. The rate of nutrient consumption is calculated at each grid based on Monod kinetics, where *R*_*i*_ is the reaction rate of nutrient *i*, *ρ* is the density of cells within the grid, Y is the biomass yield (g biomass/g CO_2_), and *σ* is the relative sucrose export activation:

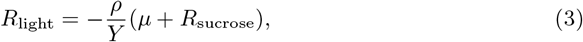

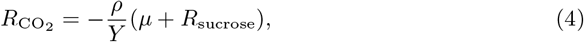

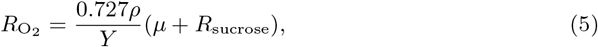

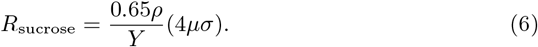

The oxygen evolution coefficient in Eq.5 was derived from the molar yield of O_2_ from CO_2_. The sucrose coefficient in Eq.6 was determined by the molar ratio of the input *vs.* output sources of carbon, which were CO_2_ and sucrose, respectively. The distribution for each soluble nutrient (*i.e.*, O_2_, CO_2_, and sucrose) in the computational domain is updated by the diffusion-reaction equation. Light attenuation through biomass was assumed to be insignificant and generally, the simulation ended before CO_2_ was depleted, so we consider the growth conditions to be overall not resource limited.

#### Algorithm 1 ABM simulation procedure

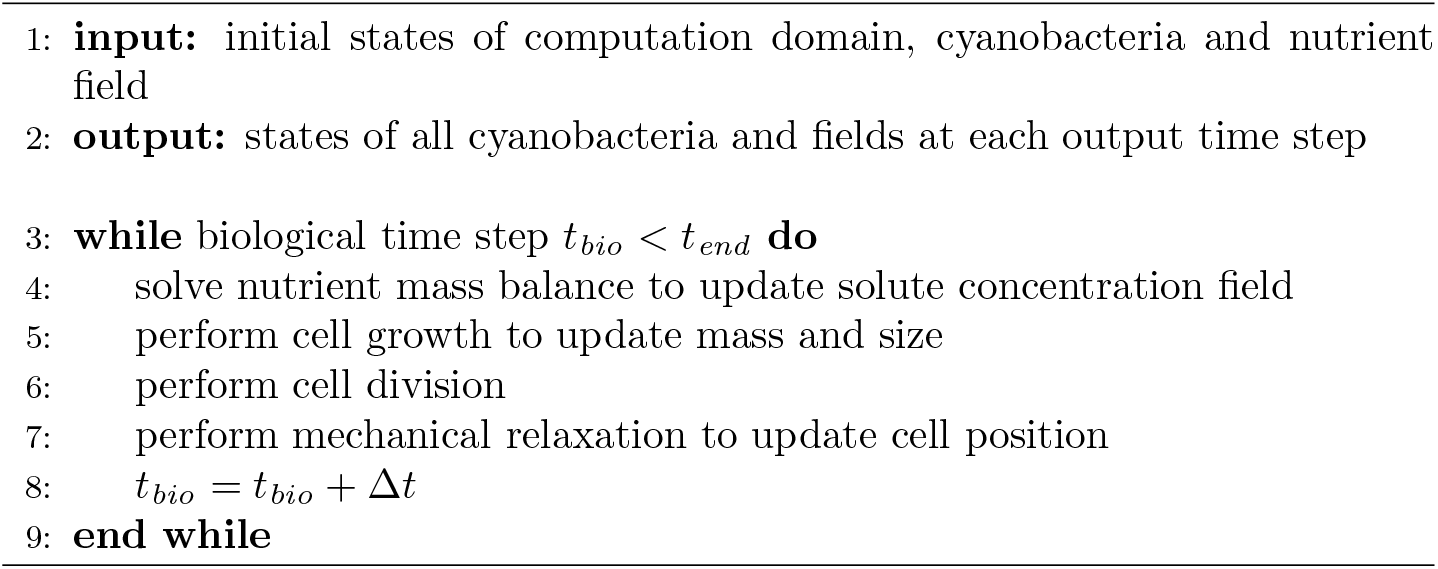

The ABM described above was realised in NUFEB [14] which is built on top of the popular LAMMPS [21] molecular dynamics software, allowing for its deployment on HPC facilities. The ABM simulation proceeds as described in Algorithm 1. The initial states of the system are defined in an input script in the form of commands and parameters. NUFEB reads those commands and parameters to undertake a simulation. During the simulation, different processes are performed sequentially, and the coupling between multiple timescales (w.r.t, biological, mechanical, and diffusion) relies on the pseudo steady-state approximation and frozen state [22]. The nutrient mass balance equation is solved based on a second-order central difference for the spatial derivatives, and a forward Euler scheme for the temporal discretisation [14]. Non-flux Neumann boundary conditions are applied for all six surfaces of the cuboid as the system is considered to be closed. Then the nutrient concentration is solved until reaching a constant flux rate, and the total flux until the next biological step is approximated.

### 2.2 Sensitivity Analysis

The method of Morris [23] was used to qualitatively understand how the outputs of our ABM depended on its inputs. This process is formally known as a Sensitivity Analysis, SA. An excellent description of the method of Morris and how to interpret it can be found in [24, 25]. Here, we briefly summarize the method to facilitate the reader understanding this section.

The method of Morris evaluates the effect that changing each input parameter has on each output. To quantify these effects, the input parameters are randomly sampled in a *k*-dimensional *p*-level grid, Ω, where *k* is the total the number of input parameters and *p* the number of different values these input parameters can assume. The sampling is performed along a chosen number of trajectories in Ω, and the effect that changing one parameter has on an output is evaluated using what is known as an *elementary effect*. These effects, one for each input parameter, are defined as follows:

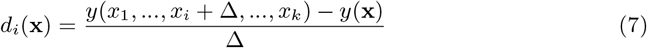

or

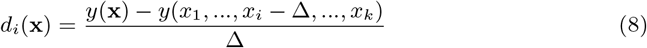

where **x** = (*x*_1_, …, *x*_*i*_, …, *x*_*k*_), *x*_*i*_ is a particular value for the *i*^*th*^ input parameter; *y*(**x**) is an output; and Δ is the shift of the *ith* parameter such that the vector (*x*_1_, …, *x*_*i*_ ± Δ …, *x*_*k*_) remains in Ω.

All the *elementary effects* of an input parameter form a distribution. The mean, *M*, and the standard deviation, *σ*, of this distribution are used to assess the effect that varying the input parameter had on the particular output. For example, if the mean *M* of *F*_*i*_ (the distribution of *elementary effects* for the *i*^*th*^ input parameter) is large, it implies that the *i*^*th*^ input parameter has a large effect on the output. If the standard deviation, *σ*, is large, it means that an *elementary effect*’s value depended strongly on where in Ω it was computed. In other words, the effect that the *i*^*th*^ input parameter has on the output is either non-linear or depends on the values acquired by the other input parameters. Following Saltelli *et al.* [25] recommendation, we also computed the mean, *M*\*, of the distribution of the absolute values of the *elementary effects*, *G*_*i*_. *M*\* is in general computed because in some cases (*e.g.* when the output depends non-monotonically on the input) *F*_*i*_ can have negative and positive terms. This can result in a small value for *M*, and be incorrectly interpreted as the *i*^*th*^ input parameter having little effect on the output.

Here the SA was used to determine the sensitivity of the population curve to the input parameters of the ABM. The population curves were generated by varying four input parameters: the initial number of cyanobacteria (*n*), the amount of *CO*_2_ (*C*), the amount of light (*L*), and the sucrose export rate (*S*). The ranges for these parameters were: [1,1000]; [15, 100]; [0.001,1]; and [0,1], for *n*, *C*, *L* and *S*, respectively. The lower limit for *CO*_2_ was found by trial-and-error: using a value below 15 led to community crashing. A total of 7500 population curves were generated using a time-step of 20 seconds. The resultant 7500 population curves were each fitted to a logistic function [26]:

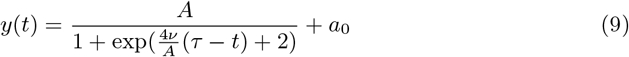

From this fitting, three output parameters were obtained for each curve: the maximum (*A*), the slope (*ν*), and the lag time (*τ*). We then used the method of Morris [23] to understand how varying *n*, *C*, *L* and *S* affected the outputs *A*, *ν*, and *τ*. Here then *k* = 4, **x** = (*n, C, L, S*), and the outputs *y*(**x**) are *A*(*n, C, L, S*), *ν*(*n, C, L, S*), and *τ* (*n, C, L, S*).

We used the method of Morris as implemented in the SALiB library [27], and fixed the number of trajectories and the plevels to 500 and 100, respectively.

### 2.3 Individual Colony Growth Metrics

Local information describing individual colony growth was extracted by analyzing images derived from simulation results. The initial colony sites were placed using one of three different spatial patterns: aggregate (attractive clustering), regular (repulsive particles), and random (2D Poisson distribution).

The initial colony locations were pre-generated in R [28] using the spatstats [29] package. Seed locations were generated with the parameters given in Table 1. Overlapping points (determined by bacterial diameter) were removed prior to simulation.

**Table 1:**
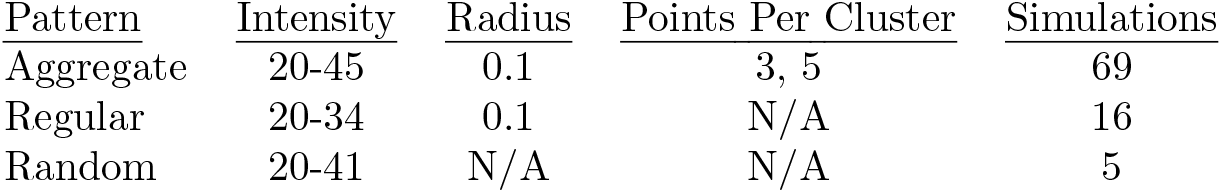
Parameters describing initial seed locations used in simulations. Intensity refers to the *realized* intensity, defined as actual number of points generated. Radius controls the area of repulsion for the regular pattern and area of attraction for the aggregate patterns. Points per cluster is the requested average value.

During each simulation, cell sizes and locations were recorded recorded every 100 timesteps. Those data were then used to generate synthetic plate images similar to those which would be observed using an optical microscope. The synthetic plate images were generated with a Python [30] script employing OpenCV [31] and Matplotlib [32]. At each timestep a two-dimensional top-down projection of the three-dimensional cell locations was produced. The initial cells were each assigned unique colony identifiers. In subsequent timesteps, new, unlabelled cells were inferred to have spawned from the closest labelled cell and assigned the appropriate colony identifier. Unlabelled cells were processed? in order of increasing distance from labelled colonies. The area of a colony at each timestep was equal to the visible area of its constituent cells in the two-dimensional projection.

Multiple approaches towards estimating colony area from the simulation data were possible, each with advantages and disadvantages. Our major criterion was to produce inferences from simulation data which may be fairly compared to those arising from actual micrographs. The approach described here was chosen because it closely resembles the way colony area is inferred from real-word light microscopy images of untagged colonies.

Each initial cell’s “room to grow” was also analyzed using Voronoi tessellation. [33] Briefly, within a group of random points, a facet associated with point *X* produced by Voronoi tessellation delineates the area closer to point *X* than all other points. Voronoi tessellation has been used in previous work [34, 35] to explore spatial effects in microbial competition. In addition to the commonly used facet area, we also recorded the number of edges for a facet (thus, the number of competing neighbors), the facet perimeter, the distance between the facet centroid and seeding location, and whether the facet is on the edge of the simulation boundary (Fig. 1).

**Figure 1:**
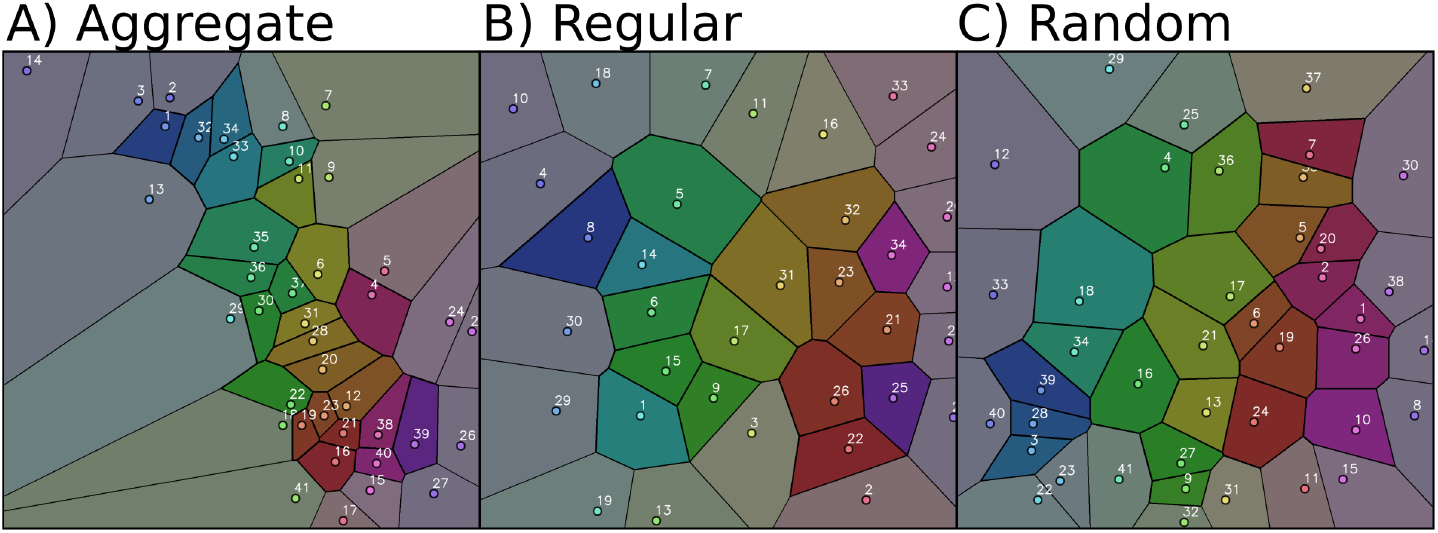
Examples of aggregate (A), regular (B), and random (C) seeding locations and their discernibly different Voronoi tessellation patterns. Regular seeding produces relatively uniform tessellation facets when compared to aggregation. Random seeding produces an intermediate tessellation; note the disparity between cells 17 and 9. Non-edge facets are highlighted with more saturated color. Intensities for the aggregate, regular, and random patterns were respectively 41, 34, and 41, and these specific simulations used to generate this figure are identified by run number: 863, 97, and 6.

The ultimate goal was to determine if spatial patterns and their associated Voronoi tessellations could be used to predict colony success, which required a defined metric quantifying “success”. Previous work [35] has measured colony success using a winner index *W I*, which is the ratio of the final area of a colony *A*_*P*_ to the area of its associated Voronoi facet *A*_*V*_ as given by Eq. 10. However, *W I* is most suitable when there is strong boundary competition and colony sizes have achieved steady-state, such as when a plate becomes fully overgrown. [35] Our simulations, however, focus on non-steady-state growth and terminated before the plate became fully overgrown. Further, we are simulating photoautotrophs which, as simulated here, do not compete for light, a major resource. Thus boundary competition was not a strong factor. To quantify colony success under those conditions, we instead used the median-normed colony area, which is the area of a specific colony *AP* normalized to the median area of all colonies for that timestep *A*_50_, as given by Eq. 11. A median-normed colony area of 0 indicates an unremarkable colony size while a positive value indicates success. A negative value, likewise, indicates a colony which grew poorly relative to others.

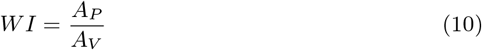

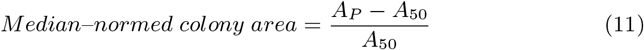

For each initial spatial pattern type, the relationship between colony success and the following factors was explored in R using both single and multiple linear regression: *facet area*, *facet perimeter*, *number of facet sides*, *distance between seed location and facet centroid*, and *facet aspect ratio*. Regressions were carried out independently on edge- and non-edge facet data. Linear regressions were carried out on both unscaled and natural log transformations of the factors and evaluated based on p-value, *R*^2^, and effect size. Multiple linear regressions without interactions were evaluated, using stepwise removal based on significance (*α* < 0.05) and Akaike Information Criterion (AIC) scores and then manually adjusted to remove terms with a variance inflation factor (VIF) greater than 2.75.

### 2.4 Microbial Growth Metamodel Based on predRNN

A metamodel based on a Recurrent Neural Networks (RNN) was trained using the data generated by our ABM model as implemented in NUFEB[14]. We chose a neural network architecture known as predRNN[15] due to its ability to capture the spatio-temporal flow of information embedded in sequences of images, *i.e.*, videos. Indeed, predRNN has been used to accurately predict future frames in videos showing numbers moving on a screen [36], and humans performing different actions [37]. Here we probe whether a metamodel based on predRNN can predict the population and colony growth of sucrose exporting cyanobacteria [3].

From the images generated in section 2.3, we created a total of 53 videos, each of 20 frames and spanning a duration of about 97 hours (timestep = 10s; total number of timesteps= 35000). This data alone was insufficient to train predRNN. Thus, as is customary in the machine/deep learning community, we augmented the data using various transformations. These transformations included rotations, translations, blurring, and noise addition, and were performed with the AtomAi package [38]. To reduce the computational cost during training, we also resized the frames from their original 300×300 size to 100×100. This pre-processing ultimately generated a total of 371 videos, each one of 20 frames.

Training predRNN was performed by dividing the set of 371 videos into training (80%) and validation sets (20%), whereas testing was performed on 17 pre-selected videos which were neither part of the training nor the validation sets. During the training process, the first 10 frames of each video were used to predict the subsequent 10. The accuracy of the predictions were then determined by comparing the predicted frames 11-20 to their “groundtruth” counterparts, *i.e.* the original frames 11-20. Comparison was performed using the metrics discussed in section 2.3, as well as the Mean Squared Error (MSE), which compares images pixelwise, and the Learned Perceptual Image Patch Similarity (LPIPS), which uses an image classification network (AlexNet) to compare layer activations between the predicted and groundtruth frames [39]. MSE and LPIPS are both routinely used in the machine/deep learning community while the other metrics are more specific to the field of microbiology. Details on the training can be found in the Supplementary Information Section.

## 3 Results

### 3.1 Sensitivity Analysis

During the SA analysis we noticed that our ABM model has two regions, with the boundary defined by the initial population size (*n*). When *n* is in the range [50, 1000], referred to from now on as region I, the population curves have smaller *A* (maximum), *ν* (slope), and *τ* (lag time), than when *n* is in the range [1, 50], region II. For this reason, we performed a Morris analysis for each region separately.

In what follows we use *M*\*’s value to rank the input parameters effect on the corresponding output. For understanding the relation between input and output, we first plotted the output *vs.* the input, and then looked at the corresponding *M*, *M*\* and *σ* values. For simplicity, we include here the results of the Morris analysis for *A* only, and present the results for *ν* and *τ* in the Supporting Information. The results for *A* are summarized in Table 2.

**Table 2:**
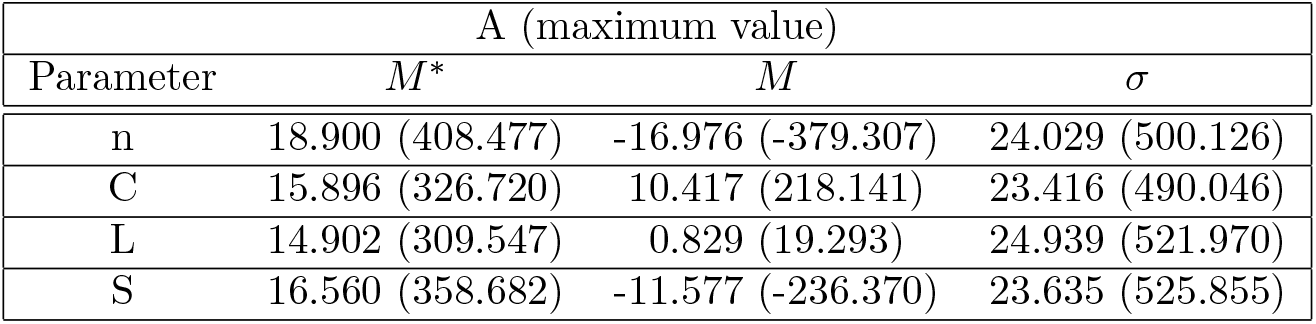
Morris analysis for *A* in regions I (II).

In both regions the input parameter that affects *A* the most is *n*, followed by *S*, *C* and *L*. The *M*\* values are strikingly different in the two regions. The reason for this becomes clear when *A* is represented *vs. n*, *C*, *S* and *L* for each region. A comparison between Figs. 2E-H and 2A-D show that *A* can reach values up to 10 times larger in region II than I, respectively. In other words, smaller initial populations lead to bigger colonies. Thus, varying an input parameter in region II can significantly affect *A*, which explains the large *M*\*. In region I (II), *A* decreases with *n* and *S*, Figs. 2A,D(E,H), and increases with *C*, Fig. 2B(F). The dependence on *n* is almost decreasing monotonic, explaining why *M* and *M*\* have similar values but different signs. *A*’s dependence on *n*, *C* and *S* makes sense: the more bacteria initially present, the more competition for resources exists, and the smaller the opportunity for growth; the more *CO*_2_ there is initially, the greater the growth; and the more sucrose is secreted, the smaller the growth. Finally, in region II the correlation effects among the input parameters is stronger, and this is captured by the large values of *σ* in this region as compared to those in region I.

**Figure 2:**
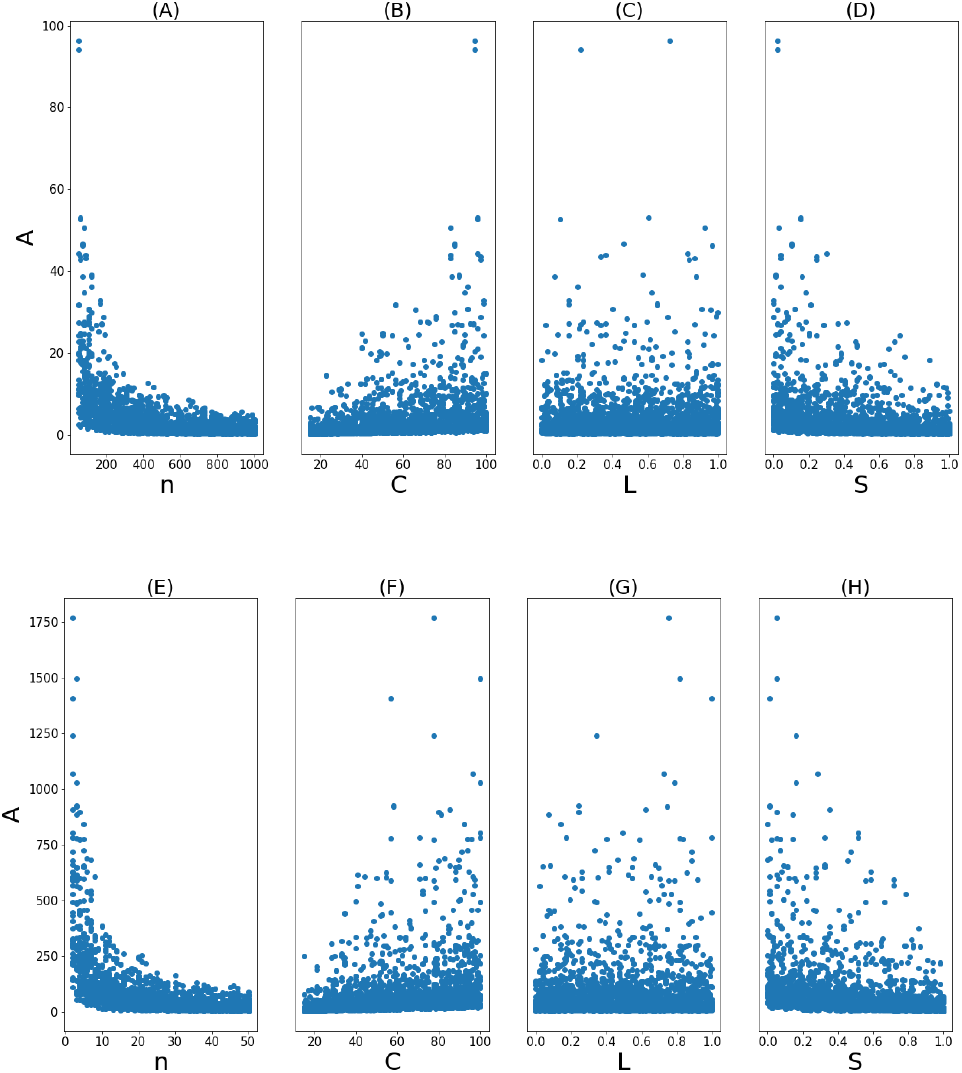
Dependence of *A* the four input parameters. Sampling for the input parameters was performed with a Morris analysis. The population curves produced were fitted to a logistic function. From this fitting we extract *A*, which stands for the maximum of the population curves. (A)-(D), results for region I; (E)-(H), results region II.

We have intentionally set aside the discussion of *L*’s effect on *A* because this case is unique. *M*\*’s and *σ*’s values are comparable to the others, but *M*’s value is significantly smaller than the rest. As for *A*’s dependence on *L*, there is not an obvious correlation, neither in region I nor II, see Figs. 2C,G, respectively. This is probably a consequence of the fact that *L*’s value remains fixed during the simulation: unlike *C*, *L*, is not a limiting factor. Thus, a population can initially grow more or less depending on *L*’s value, but then it’s *C*’s availability what determines how much and for how long this population will grow.

### 3.2 Colony success and Initial Spatial Distribution

As expected, the three seeding spatial patterns produced Voronoi tessellations with differently distributed facet areas (Fig. 3A). Aggregation produces clustered seeding locations which results in many smaller Voronoi facets, skewing the distribution. In contrast, the repulsive forces present during regular spatial patterning produces results in many similar-sized facet and a centered distribution. A randomly distributed spatial pattern, which has neither attractive nor repulsive forces, results in a somewhat intermediate distribution.

**Figure 3:**
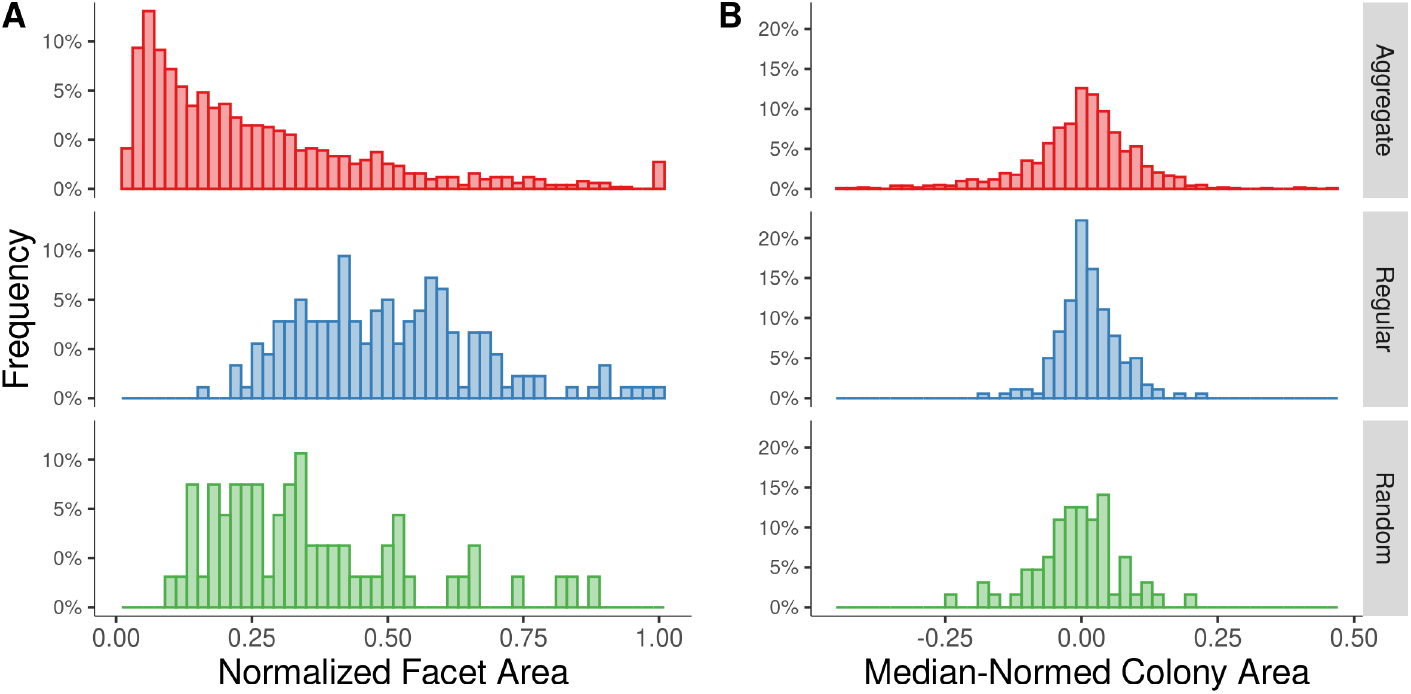
Spatial patterns of initial seed locations produce differing distributions of Voronoi facet (A) and colony (B) areas. Normalization was carried out on a per-simulation basis. Facet areas are normalized to the largest facet area in the relevant run. Colony areas were normalized to the per-run median colony area, as defined in section 2.3, Eq. 11. Edge facets were not included in this figure, but produce qualitatively similar distributions despite edge effects. Histograms including edge facets and also other factors (*e.g.* facet sides and centroid-seed distance) and are included as supplementary information.

Colony size distributions, measured as the median-normed colony area, also differed based on spatial patterning. Aggregate patterning produced a heavier-tailed distribution with higher variance. Regular patterning, as expected, resulted in sharply centered colony areas with low variance. As with facet areas, random patterning produced a colony area distribution with aspects intermediate to those of the aggregate and regular patterns.

Spatial patterning affects both facet and colony areas. However, under the weak boundary competition conditions of this study, there’s no easily visible correlation between the facet area and the colony area (Fig. 4) for non-edge colonies. Other factors, including non-facet aspects such as the log-inverse square distance between seeds, similarly exhibit a similar lack of definite correlation. On the other hand, edge-effects do appear to affect correlation under any pattern, particularly for small facets. Thus, it is important that any study correlating spatial position with colony growth must account for edge-effects.

**Figure 4:**
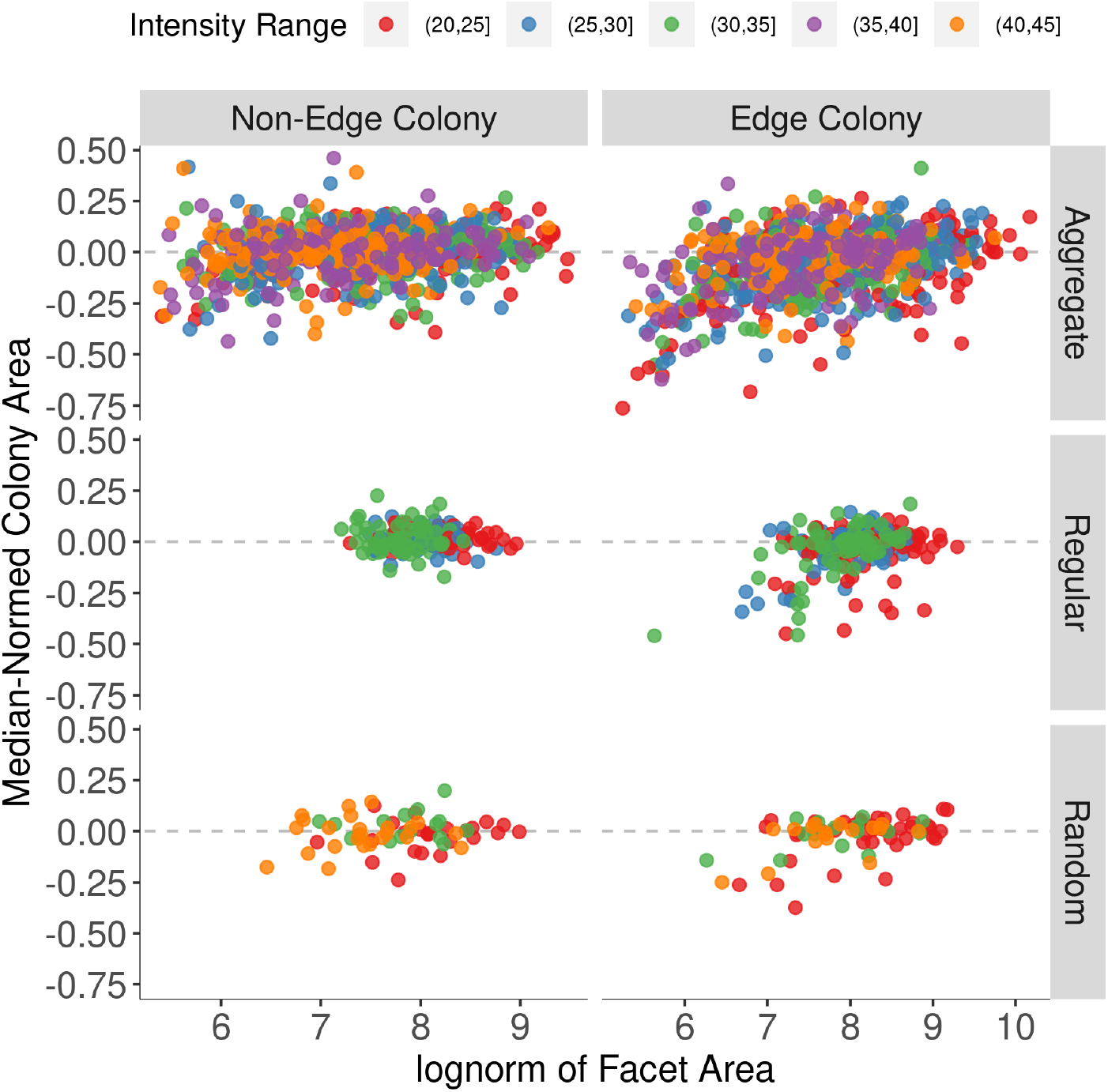
Apart from edge-effects, Voronoi facet area (log transformed) is not appreciably correlated with colony size under weak boundary competition.

Regressions on either single factors or the log transform of those factors (Table 3.2) quantitatively support the qualitative conclusions drawn from (Fig. 4). For non-edge facets, no factor significantly predicted colony success for either regular or random spatial patterning. In the case of aggregate patterning, some significant fits occurred, but most of the variance in the data (≥ 95%) remains unexplained. We believe that the major difference between aggregate patterning and the others is that the close clustering associated with aggregation leads to spatially localized strong boundary-competition situations in which Voronoi facets have been previously shown to be useful predictors of colony success.[35, 34]

**Table 3:**
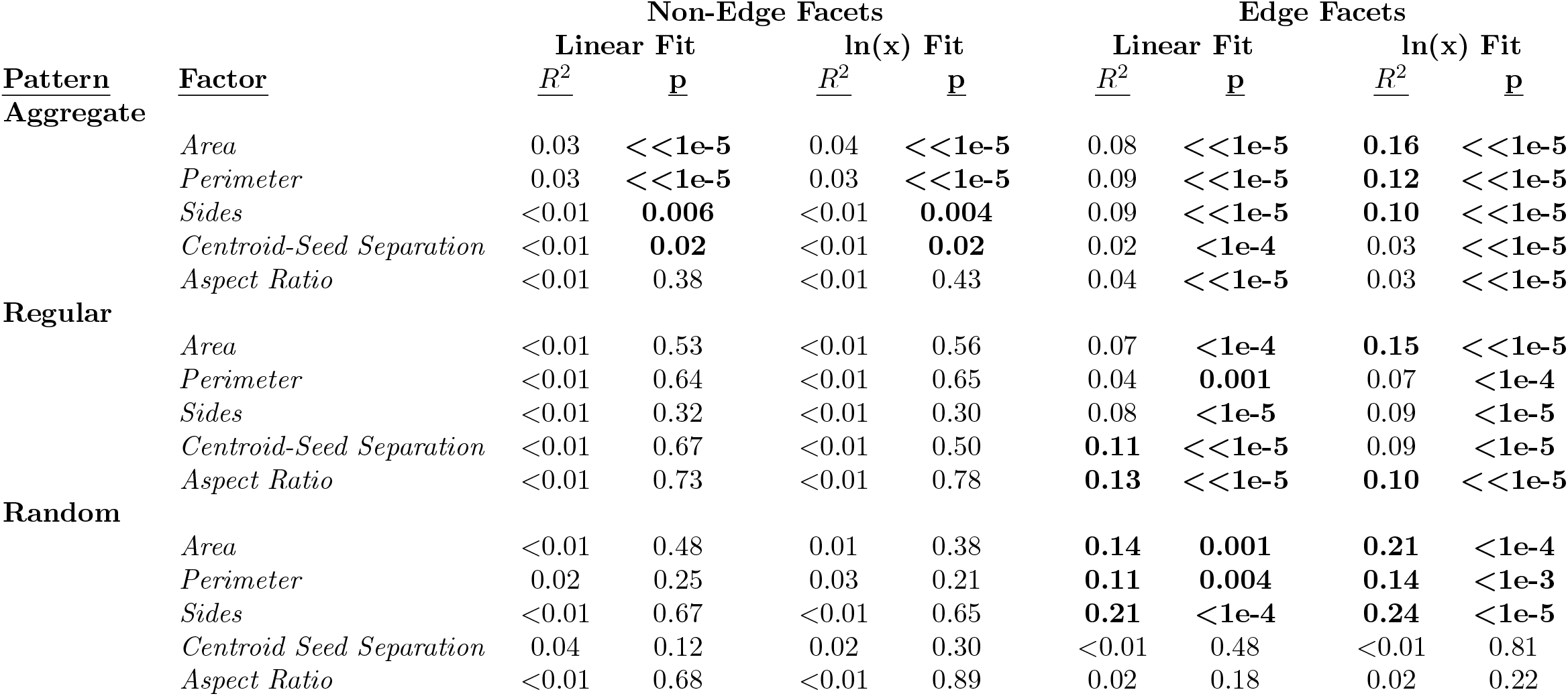
Few potential single-variable linear regressions produce useful predictions of colony size, once edge-effects are accounted for. Presented here are the fit metrics (adjusted *R*^2^ and p-value for regressions using each potential predictor variable (or its log-transformation), organized by edge- vs. non-edge facets and the spatial pattern used to place initial seed sites. Boldface highlights significance (p-value greater than 0.5) or fits which explain more than 5% variance.

| <u>Pattern</u> | <u>Factor</u> | Non-Edge Facets |  |  |  | Edge Facets |  |  |  |
| --- | --- | --- | --- | --- | --- | --- | --- | --- | --- |
|  |  | Linear Fit |  | ln(x) Fit |  | Linear Fit |  | ln(x) Fit |  |
|  |  | <u>$R^2$</u> | <u>p</u> | <u>$R^2$</u> | <u>p</u> | <u>$R^2$</u> | <u>p</u> | <u>$R^2$</u> | <u>p</u> |
| **Aggregate** | *Area* | 0.03 | **<<1e-5** | 0.04 | **<<1e-5** | 0.08 | **<<1e-5** | **0.16** | **<<1e-5** |
|  | *Perimeter* | 0.03 | **<<1e-5** | 0.03 | **<<1e-5** | 0.09 | **<<1e-5** | **0.12** | **<<1e-5** |
|  | *Sides* | <0.01 | **0.006** | <0.01 | **0.004** | 0.09 | **<<1e-5** | **0.10** | **<<1e-5** |
|  | *Centroid-Seed Separation* | <0.01 | **0.02** | <0.01 | **0.02** | 0.02 | **<1e-4** | 0.03 | **<<1e-5** |
|  | *Aspect Ratio* | <0.01 | 0.38 | <0.01 | 0.43 | 0.04 | **<<1e-5** | 0.03 | **<<1e-5** |
| **Regular** | *Area* | <0.01 | 0.53 | <0.01 | 0.56 | 0.07 | **<1e-4** | **0.15** | **<<1e-5** |
|  | *Perimeter* | <0.01 | 0.64 | <0.01 | 0.65 | 0.04 | **0.001** | 0.07 | **<1e-4** |
|  | *Sides* | <0.01 | 0.32 | <0.01 | 0.30 | 0.08 | **<1e-5** | 0.09 | **<1e-5** |
|  | *Centroid-Seed Separation* | <0.01 | 0.67 | <0.01 | 0.50 | **0.11** | **<<1e-5** | 0.09 | **<1e-5** |
|  | *Aspect Ratio* | <0.01 | 0.73 | <0.01 | 0.78 | **0.13** | **<<1e-5** | **0.10** | **<<1e-5** |
| **Random** | *Area* | <0.01 | 0.48 | 0.01 | 0.38 | **0.14** | **0.001** | **0.21** | **<1e-4** |
|  | *Perimeter* | 0.02 | 0.25 | 0.03 | 0.21 | **0.11** | **0.004** | **0.14** | **<1e-3** |
|  | *Sides* | <0.01 | 0.67 | <0.01 | 0.65 | **0.21** | **<1e-4** | **0.24** | **<1e-5** |
|  | *Centroid Seed Separation* | 0.04 | 0.12 | 0.02 | 0.30 | <0.01 | 0.48 | <0.01 | 0.81 |
|  | *Aspect Ratio* | <0.01 | 0.68 | <0.01 | 0.89 | 0.02 | 0.18 | 0.02 | 0.22 |

Edge-facets, unlike non-edge facets, were associated with regression models which did significantly predict colony success for all spatial patterns, particular regressions using log-transformed predictor variables. Although the percentage of variance, measured as *R*^2^, explains remains modest (at most 24%), it is still notably higher than the *R*^2^ of the few significant non-edge facet regressions. Others have also noted the effect of edge-distance on such correlations [40] and while we may speculate on the cause, the main import of this difference is to support our previous assertion that edge effects matter; any study correlating spatial position with colony growth must account for edge-effects to avoid spurious correlations.

Multiple linear regressions produced numerically similar and effectively low-utility (*e.g.*, low percent variance explained) models. The general predictive trends (edge-effects, low predictive power for regular and random spatial patterns, slight advantage of log transformation) remained the same as those observed using single variable regressions.

In short, under conditions in which boundary competition is minimal, classical regression techniques do not produce models which appreciably predict colony growth and success based on Voronoi tessellations and other factors derived from the initial seed sites. This is unlike the case where moderate [34, 40] or intense [35] boundary competition was present and in which Voronoi facets were useful predictors of colony growth and success.

Our results generally agree with the portions of other studies [34, 40] in which colony sizes had not yet reached steady state (as in our study) and boundary competition was presumably low. Further, we believe the lower correlation coefficients in our study, relative to those portions, can be explained by the the degree to which edge-associated colonies were removed from analysis.

### 3.3 Predicting Microbial Growth with a predRNN Meta-model

To investigate whether predRNN could be used as a metamodel of the ABM, we considered the growth of 6 different populations, each representing aggregate, regular, and random initial conditions. The ABM generated videos for each populations contained 20 equally spaced image frames, the first 10 of which were given as input to predRNN, which then returned 10 frames of predicted images. A sample comparison of the ABM simulated images and the predicted ones is shown in Figure 5. Overall, predRNN maintained the colony locations and general growths, but also exhibited small bleeding in both shape and color.

**Figure 5:**
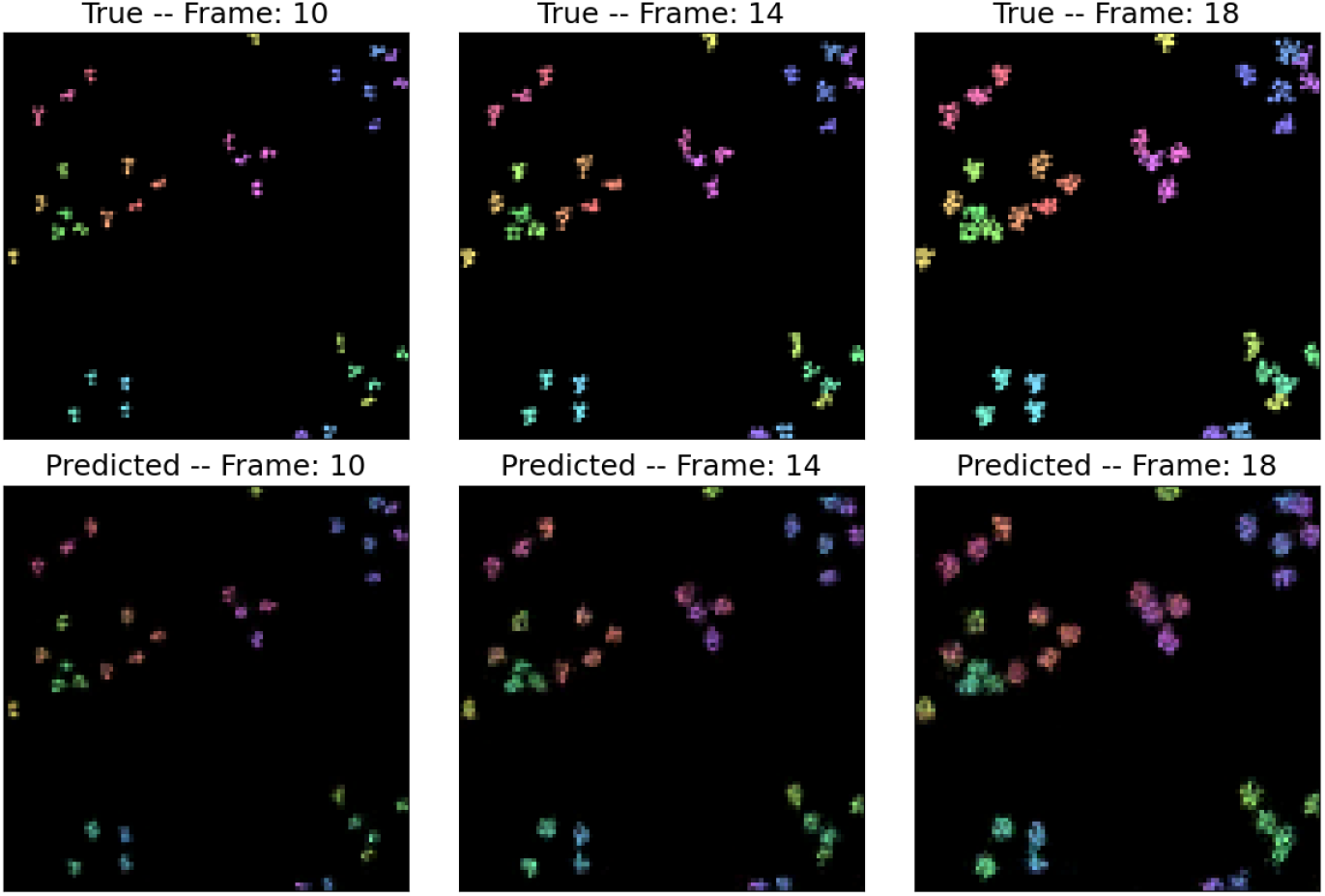
Simulation and predicted images of colony growth match well in size and distribution. Shown here is a comparison of these image outputs. Predictions begin at frame 10 in each 20 frame set.

The predicted frames 11-20 were compared to the groundtruth ones, *i.e.* the simulated frames 11-20, using MSE and LPIPS, and the metrics of section 2.3. The MSE and LPIPS values per frame are shown Figs. 7a,b, respectively. MSE and LPIPS increase with time, indicating that the accuracy of the predictions decreases with time (which is common for RNNs).

For any of the considered initial conditions, the metamodel was able to reproduce a population curve and growth rate nearly identical to that of the simulation as seen in Fig. 6.

**Figure 6:**
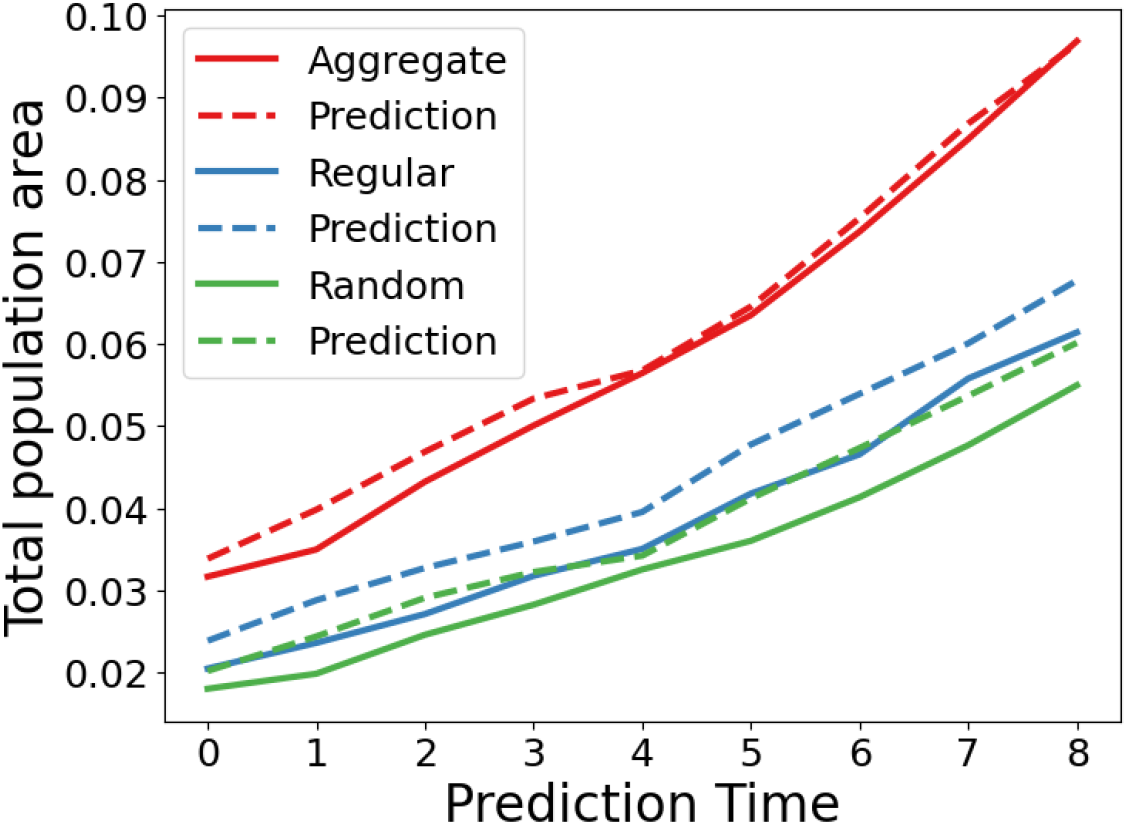
The population growth observed in the predicted images match closely those observed in the simulation. Here is shown the populations of the simulation compared with the populations

**Figure 7:**
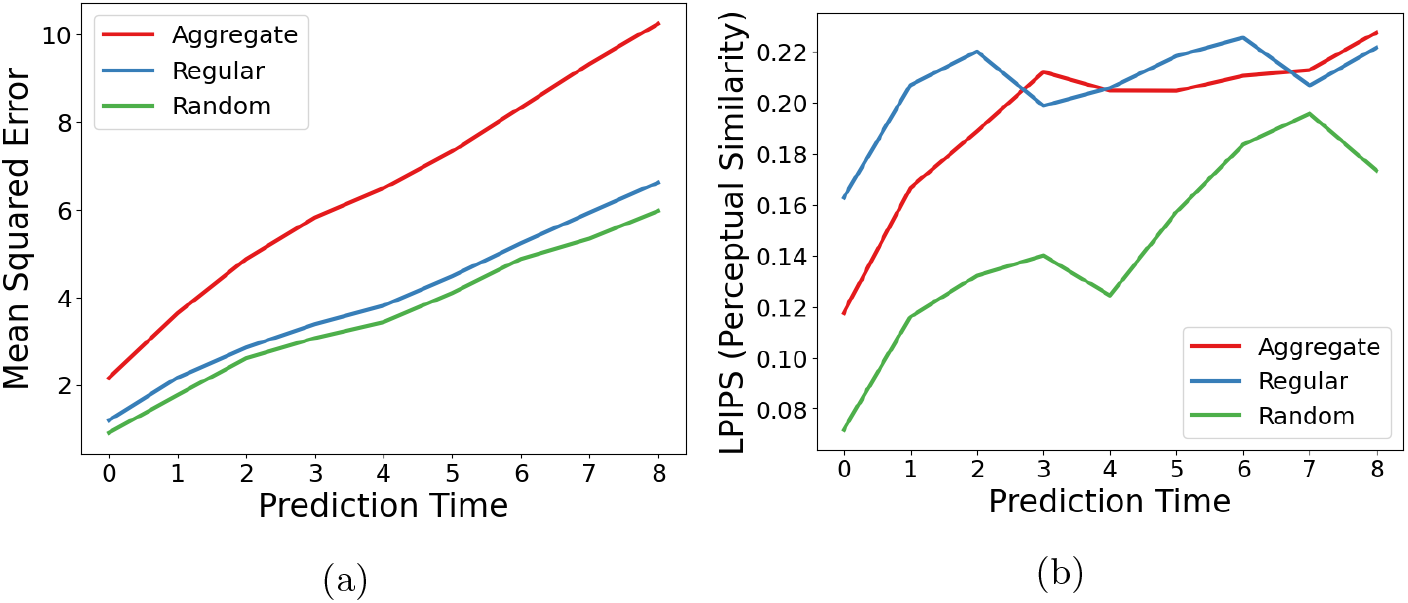
As the prediction progresses, the accuracy of the predictions declines. a) Mean squared error and b) LPIPS perceptual similarity of ABM simulated image compared to predicted image for varied initial conditions. Note that the “aggregate” case has overall higher population, resulting in larger MSE values. However, the rate of increase in MSE for all three cases is consistent.

Next, we extracted the median-normed colony area from both the groundtruth and predicted images, and compared the results in Fig. 8. It’s clear that for any of the considered initial conditions, the metamodel was able to reproduce a population curve and growth rate nearly identical to that of the simulation. Additionally, the histograms of the individual colony sizes (in median-normed colony area) also reflected well the ABM values, as seen in Figure 8.

**Figure 8:**
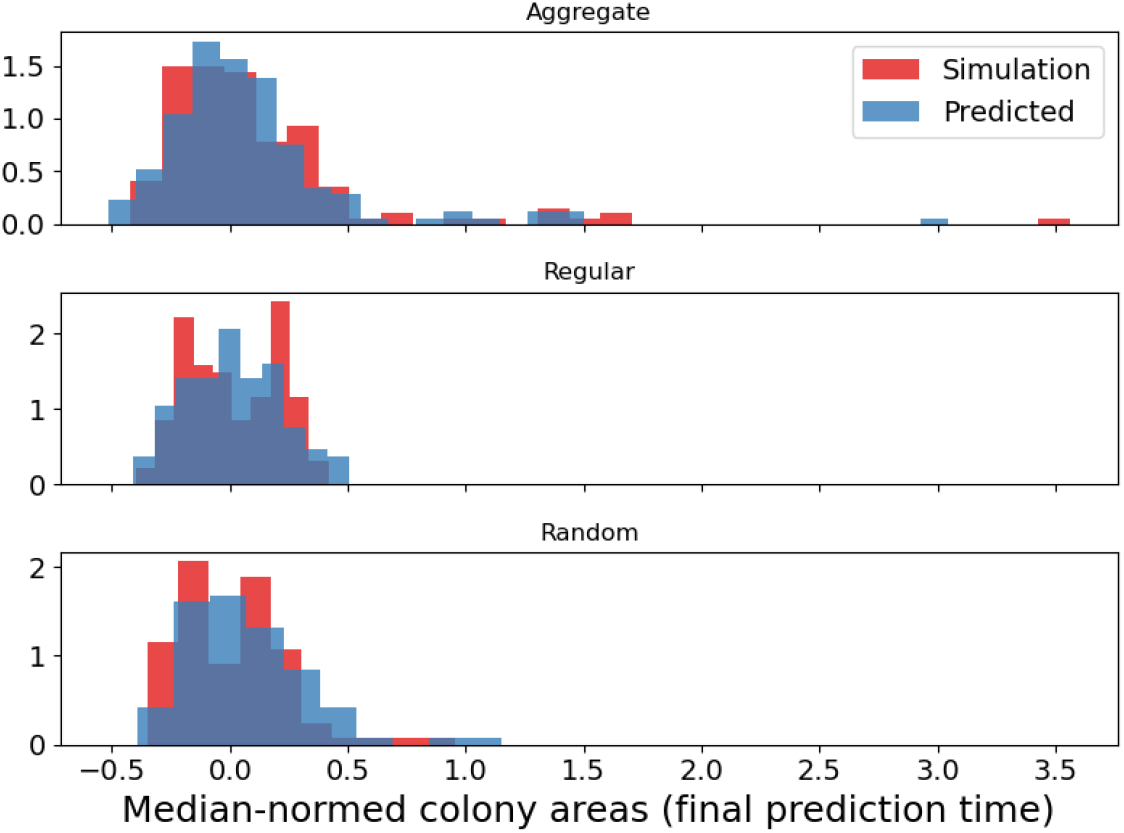
Comparison of the distribution of predicted median-normalized colony areas with the simulation median-normalized colony areas for varied initial conditions.

This demonstrates that, unlike classical correlation, the metamodel was capable of predicting large and small scale population dynamics despite the transient growth conditions and weak boundary competition.

Finally, an important observation that resulted from the analysis in section 3.2, is the identification of the most successful individual colonies using the median-normed colony area. This metric requires not only predictive accuracy on averaged measures, such as the area histogram, but also local accuracy for individual colony growth rates. Figure 9 demonstrates that although the prediction is not exact in identifying the relative colony sizes, it holds strongly in correlation: the colonies that exhibited relatively large areas in simulation were also relatively large in the prediction. Although this is an important validation, the metamodel does receive as input information from the first half of the simulation which indicates promising colonies. However, it is encouraging that the metamodel can propogate this information further into the simulation and opens the door to more intricate predictions for systems that grow to a full lawn.

**Figure 9:**
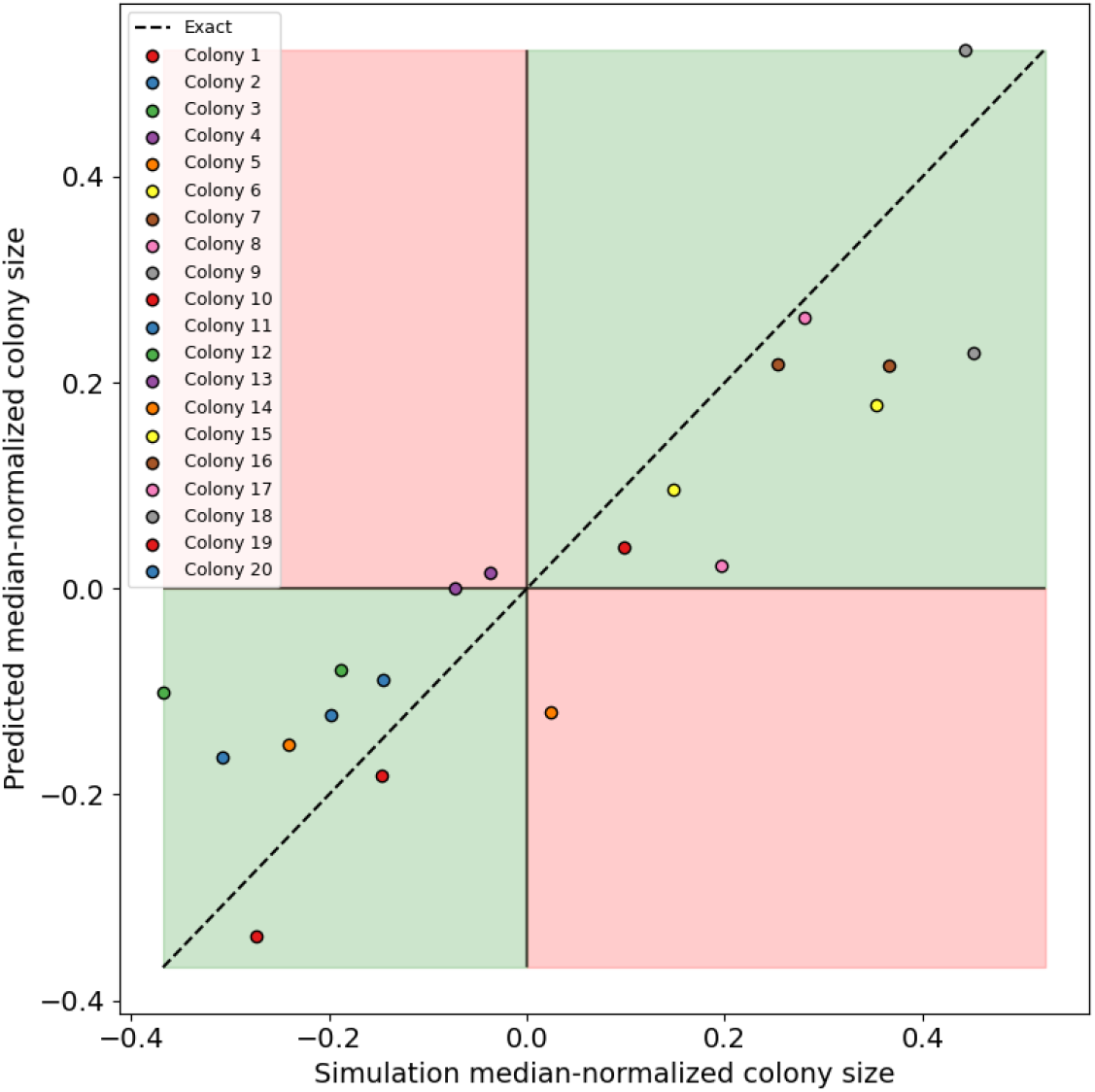
Aggregate colony comparison of median-normalized colony area between prediction and simulation at final timestep of prediction. The green areas highlight areas in which the simulation and prediction are correlated (large colonies in simulation are large in prediction) and the red areas where they are not correlated.

These results demonstrate the feasibility of accelerating or extending ABM simulation results via recurrent neural networks such as PredRNN. Further information on the time required for training and prediction using PredRNN can be found in Appendix **??**.

## 4 Discussion

We have developed a computational framework to investigate the growth of a engineered strain of *Synechococcus elongatus* PCC 7942 that exports large quantities of sucrose. The framework combines high performance ABM simulations with statistical analysis and deep learning techniques. The goal was to pave the road towards developing a simulation framework to aid pure wet-lab experiments of microbial populations by exploring the solution space *in silico* first. To this end, we needed to ensure that the framework was efficient and accurate. Efficiency was ensured by implementing the ABM into NUFEB [14], which uses LAMMPS [21], a highly parallelizable computational framework. Accuracy can only be evaluated by comparing directly with experiment. In the absence of it, we investigated how growth in our simulations depended on nutrients and the initial distribution of colonies. This information provided an understanding of the inner workings of our ABM, and will facilitate future modifications, if they were to be needed. The ABM accounted for nutrients intake, as given by *CO*_2_ and light, and sucrose secretion. We found that growth proceeded differently depending on whether the initial number of bacteria was smaller or larger than 50. Below 50, large populations were generated and stronger correlations among the input parameters were observed. A counter-intuitive result was observed when the dependence of growth on light was analyzed. This dependence did not seem obvious, and this is a probably due to the fact that light is not a limiting factor in our simulations. As a consequence, the effects that it might have on growth are washed out by the effects caused by limiting factors, specifically *CO*_2_: once *CO*_2_ diminishes below a certain threshold, growth stops, no matter how much light is in the sample. Keeping light constant during the simulation is probably not optimal, yet it’s not obvious to describe light changes across a sample and over time with an ABM. As a first approximation, we chose to keep light constant. Future comparison with experimental data will determine if this assumption holds.

Three different initial colony distribution styles (aggregated, regular, and random) were generated to investigate the relation between colony growth success and seeding locations. Although the different seeding patterns produced, as expected, differing ‘local neighborhoods’ (measured as Voronoi facets), the ultimate success of the colonies did not strongly correlate to any aspect of the neighborhood (*e.g.* facet area, perimeter). This is in contrast to previously reported findings [34, 35, 40] and we attribute this difference both to the low amount of spatial competition in our simulations and the removal of confounding edge-effects.

Finally, we demonstrated that a metamodel based on a RNN is capable of reproducing ABM simulations at a lower computation cost. It’s not the first time that an artificial neural network is used to build a metamodel of an ABM. For example, in [41], we showed that a fully dense neural network was capable of reproducing the ABM population curves for *Pantoea*. What is novel about using a metamodel based on a RNN is the fact that with such a metamodel one can reproduce ABM synthetic images of growth. From these, one can extract not only population curves, of course, but also colony metrics that could reveal growth patterns. It’s in this last aspect where RNN based metamodels become specially useful: if the experimental system size becomes unattainable with ABM, a RNN metamodel could take over and make predictions which accuracy will be determined by comparing *directly* with experimental images of growth, not just with population curves. This brings simulations in microbiology a step closer to Pattern Oriented Modeling [13].

## Data Availability statement

The code, smaller datasets, and instructions for reproduction have been place in a git repository at https://github.com/Jsakkos/nufeb-cyano-rnn. Larger datasets have been made available as part of an Open Science Foundation (OSF) repository at https://osf.io/fmwbp/.

## Acknowledgments

This research was conducted at the Center for Nanophase Materials Sciences, which is a DOE Office of Science User Facility. This research used resources of the Compute and Data Environment for Science (CADES) at the Oak Ridge National Laboratory, which is supported by the Office of Science of the U.S. Department of Energy under Contract No. DE-AC05-00OR22725. J.W. was supported by a U.S. National Science Foundation fellowship under Award No. 2007151. This material is also based upon work supported by the U.S. Department of Energy, Office of Science, Office of Workforce Development for Teachers and Scientists, Office of Science Graduate Student Research (SCGSR) program. The SCGSR program is administered by the Oak Ridge Institute for Science and Education (ORISE) for the DOE. ORISE is managed by ORAU under contract number DE-SC0014664. All opinions expressed in this paper are the author’s and do not necessarily reflect the policies and views of DOE, ORAU, or ORISE.

## Licenses and Permissions

This manuscript has been authored by UT-Battelle, LLC under Contract No. DE-AC05-00OR22725 with the U.S. Department of Energy. The United States Government retains and the publisher, by accepting the article for publication, acknowledges that the United States Government retains a non-exclusive, paid-up, irrevocable, world-wide license to publish or reproduce the published form of this manuscript, or allow others to do so, for United States Government purposes. The Department of Energy will provide public access to these results of federally sponsored research in accordance with the DOE Public Access Plan (http://energy.gov/downloads/doe-public-access-plan).

